# Demographic analysis of Israeli *Carpobrotus* populations: management strategies and future directions

**DOI:** 10.1101/2020.12.08.415174

**Authors:** Ana Bogdan, Sam C. Levin, Roberto Salguero-Gómez, Tiffany M. Knight

## Abstract

*Carpobrotus* species are harmful invaders to coastal areas throughout the world, particularly in Mediterranean habitats. Demographic models are ideally suited to identify and understand population processes and stages in the life cycle of the species that could be most effectively targeted with management. However, parameterizing these models has been limited by the difficulty in accessing the cliff-side locations where its populations are typically found, as well as accurately measuring the growth and spread individuals, which form large, dense mats. This study uses small unmanned aerial vehicles (UAVs, drones) to collect demographic data and parameterize an Integral Projection Model of an Israeli *Carpobrotus* population. We validated our data set with ground targets of known size. Through the analysis of asymptotic growth rates and population sensitivities and elasticities, we demonstrate that the population at the study site is demographically stable, and that reducing the survival and growth of the largest individuals would have the greatest effect on reducing overall population growth rate. Our results provide a first evaluation of the demography of *Carpobrotus,* a species of conservation and economic concern, and provide the first stage-based population model of a representative of the Aizoaceae family, thus contributing to our global knowledge on plant population dynamics. In addition, we demonstrate the advantages of using drones for collecting demographic data in understudied habitats such as coastal ecosystems.

## Introduction

Non-native invasive species are widely recognized as a major threat to native biodiversity, ecosystem function, and economic interests at a global scale [1,2]. The succulent/sub shrub species of the *Carpobrotus* genus are particularly problematic as they are “transformers” of frail coastal habitats, forming mats that prevent sand movement and disrupt normal dune processes of disturbance and succession [3–5]. Most *Carpobrotus* species are native to the Cape region of South Africa, and some notorious species (e.g., *C. edulis, C. acinaciformis)* are now naturalized in Mediterranean climate regions on all continents except Antarctica [6]. In Europe, conservation areas have dedicated ~1,000,000 EUR/year to controlling *Carpobrotus* invasions [6–9].

*Carpobrotus* species decrease native plant diversity in the places they invade by direct and indirect competition with the local flora [4,6,10–12]. Individuals of the *Carpobrotus* genus compete directly with native plants for space, water, and nutrients [13,14], and indirectly by changing the soil pH [4], salt content, moisture level, nutrient content and microbial activity [6]. *Carpobrotus* litter disrupts natural soil nutrient cycles, increasing nitrogen and organic matter content and releasing allelochemicals that hinder seed germination and root growth of some native plants [15,16]. These disruptions to biotic and abiotic processes can hinder management efforts as they can persist for years after the invasion has been eradicated [17,18].

Structured population models (e.g., matrix population models [19], integral projection models [20]) of invasive plants provide a tool for generating comprehensive fitness estimates and quantifying relative importance of vital rates (e.g. survival, growth). With these estimates, and an explicit consideration of management costs [21,22], managers can optimize management strategies to limit population growth [19,23–25]. To date, most research on *Carpobrotus* has focused on a determining its effects on other species in their ecological communities and experimentally assessing individual vital rates for vulnerability to various management strategies [16,26–28]. Succulents are an exceptionally diverse plant group, both functionally and phylogenetically, and generate both positive (e.g. farmed agaves, pineapples) and negative economic impacts (e.g. invasive species like *Carpobrotus spp, Lampranthus spp,* and *Opuntia spp,* [29]). However, no studies have conducted a comprehensive demographic analysis of any species in the *Carpobrotus* genus. In addition, to our knowledge, stage-structured demographic studies on any member of the Aizoaceae family are non-existent, and most of the information we have on the population dynamics of succulents comes from the Cactaceae family [30].

Structured population models are scarce for plants in coastal habitats [30]. Plants in these habitats provide vital ecosystem services such as acting as a buffer against coastal hazards (wind erosion, tidal inundation during storm or hurricanes and wave overtopping), as well as promoters of biodiversity by providing travel corridors and a wide array of habitats for endemic plant, birds, reptiles and invertebrates [31,32]. In addition, coastal plants stimulate dune growth by trapping and stabilizing wind-moving sand and a vegetated dune system will have sufficient sand to restore beaches to their original state after a storm [33]. Given the importance of these vulnerable coastal communities, we need further research to understand the demographic processes of species in their communities in order to preserve them. In general, demography is a data-hungry and time-consuming enterprise in any habitat [34]. However, demographic data are particularly challenging to collect in coastal populations, which typically occur on inaccessible terrain, such as coastal bluffs [35]. Only a few prior studies have managed to quantify demographic variables safely and precisely in such challenging environments (e.g., freehand climbing in [36], targeting individual plants that can be reached without climbing equipment, [37]). The challenge of difficult terrain can be overcome by using unmanned aerial vehicles (UAVs), a rather recent technological advance in ecological monitoring studies [38–42]. UAVs allow sampling of coastal bluffs, which can otherwise be difficult and dangerous to reach [42,43]. UAVs provide high resolution imaging [39,41,42,44], and require little space for landing/take-off and minimal piloting expertise [39,40,43]. In addition, data collected from UAVs have been shown to be less error-prone that those collected directly by humans [45].

Here, we quantify the demography of a *Carpobrotus* population using small unmanned aerial vehicles (UAVs) to collect data and construct Integral Projection Models (IPMs). We project the asymptotic population growth rate (*λ*) and calculate sensitivities and elasticities of *λ* to determine which vital rates (survival, growth, and reproductive success as both flowering probability and number of flowers) have the greatest effect on population growth rate if altered by management. We found that the size of *Carpobrotus* individuals predict its vital rates and that its demography is typical of a long-lived polycarpic plant. We demonstrated that drones allow for collection of demographic data in previously inaccessible locations, thus showing their great promise for filling global research gaps. We discuss the consequences of our results for future *Carpobrotus* control actions and emphasize the need to target survival and growth of large plants.

## Methods

The study was conducted between April 2018 and April 2019 on a cliff top and cliff side overlooking the Mediterranean Sea, just north of Havatselet Ha’Sharon in Israel (32.36403 N, 34.8578 E, approx. elevation 25 m a.s.l., Figure 1). The area (hereby termed Havatselet) is characterized by heavy recreational use and is occasionally subject to small landslides from the cliff top down to the beach at the bottom. *Carpobrotus* is widespread in gardens in the surrounding area, and it is likely that this population escaped cultivation and established itself. We could not find any information on the age of invasion, but the size and consistent spatial distribution of the individuals suggests that it is a mature population.

**Figure 1:**
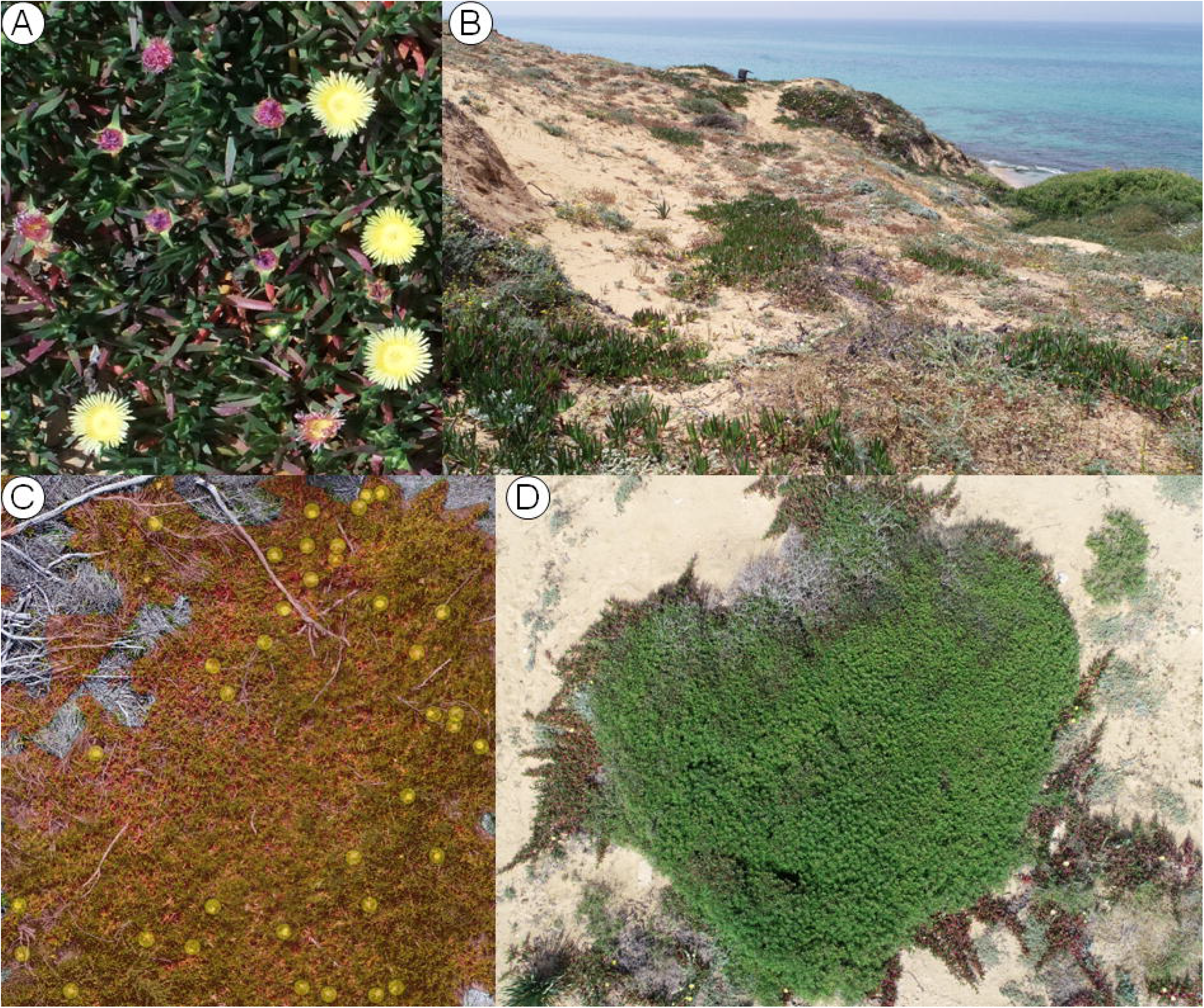
A close up picture of the *Carpobrotus spp.* present at the Havatselet field site (a), a wider angle shot of part of the site (b), polygons overlaid on a *Carpobrotus spp.* individual after orthomosaicing, as well as a point layer used to count flowers (c), and single nadir image from the drone (d). 1695 of these were calibrated and used to generate the 2018 orthomosaic, and 1021 of these were calibrated and used to generate the 2019 orthomosiac.

*Carpobrotus* species are known to hybridize with each other, raising numerous identification constraints [46,47]. Without genetic analysis, we were unable to determine exactly which species we worked with at Havatselet, but based on reports of species presence in Israel [5,48] it is safe to assume that most probably the species were *Carpobrotus edulis, C. acinaciformis* and *C. chilensis* together with their hybrids, these being also the most problematic *Carpobrotus* species [15,46,49]. Hybrid *Carpobrotus spp* can have unique behaviors in their invaded range, as hybridization contribute to evolutionary changes in the dynamics of invasion [50], meaning our results may not apply to hybrids on other continents.

We developed a flight plan to map the population using DJI Ground Station Pro v2 [51] for iPad. This application allows users to generate polygons over areas of interest and computes optimal flight paths for surveying given a desired set of photo overlap, map resolution, and cameras on the drone itself. We selected a flight path that generated a resolution of ~0.35cm/pixel with ~85% side and front photo overlap, with the camera pointed 90 degrees downward (i.e. nadir imagery). The average width of open flowers is 7.22 cm (standard error: 0.042 cm) and unripe fruits was 2.2 cm (standard error: 0.014 cm) at the Havatselet population (SC Levin, unpublished data), so this resolution is sufficient to allow us to mark the majority of both in the resulting maps (orthomosaics, see below for more details). DJI Ground Station Pro only offers an option to fly at a specific altitude above the takeoff point, rather than keeping a fixed altitude over the terrain. Thus, the flight was conducted by hand so that we could maintain a consistent altitude over the variable terrain. Transects were flown with a DJI Phantom 4 Pro v1 [51] along the lines of the flight plan generated by DJI GSP and images were recorded at 1.8 – 2m intervals. When batteries reached a critical level of power (i.e. 20%), we landed the drone, switched the batteries out, and the flight was resumed from the last stopping point. The estimated area covered was 2.063 hectares. The mission took approximately 1 h 40 min from 12:50 PM to 2.30 PM and five battery charges were necessary.

Once all images were captured in 2018, individual photos were processed into a single composite, georeferenced orthomosaic map using Pix4Dmapper [52]. We drew individual polygons around contiguous *Carpobrotus* plants (ramets), assigned each a unique ID number, and counted flowers using a point layer in QGIS 3.4 [53]. We repeated the process in 2019 by overlaying the 2019 map on the 2018 map and marking polygons for all surviving individuals as well as new recruits also in QGIS. Individual plants that were not found again in 2019 were assumed dead. Maps were aligned between April 2018 and 2019 using the Georeferencer GDAL plugin [54] in QGIS because the georeferencing of the orthomosaics was only accurate to ~5m. This plugin allows users to manually find and mark common points on both images and set them as reference points for transformation between the old coordinate system (map coordinates in 2019) and the new coordinate system (map coordinates in 2018). Pix4D generated multiple blocks of images during the orthomosaic optimization step in 2018, but not 2019, resulting in slight warping of the final image in 2018. We used the thin plate splines transformation for the coordinate systems and nearest neighbor resampling method. The reference points are available in the supplementary materials. In addition to drawing polygons around the individual plants, we drew polygons around targets of known sizes and computed the ratio of calculated areas under the polygon vs the known sizes. We then re-scaled all computed sizes of plants using this ratio. Based on the ground truthing procedure, ramet sizes were underestimated by ~12% using the polygons from QGIS. Thus, all measures of size were re-computed before log transformation. The technique of overlaying maps from subsequent years adequately captured populations at resolutions high enough to identify plants and flowers (Online Supplementary Materials). We identified 276 distinct ramets in 2018, re-found 233 in 2019, and were able to mark 10 new individuals in 2019. Georeferencing accuracy varied throughout the orthomosaic due to the blocks generated in 2018, but was never off by more than 4m, and we are confident that we correctly identified each ramet in 2019 as either existing or new.

To model vital rates, we considered how an individual’s size (*z*) influences its survival, growth, and reproduction. We constructed a series of progressively more complicated generalized linear models for survival (*s*(*z*)) and growth to size *z*’ conditional on survival (*g*(*z‘,z*)) using an intercept only, log-transformed surface area, and a 2^nd^-order polynomial of log-transformed surface area as an explanatory variable. Additionally, we fit generalized additive models with 8 knots and 6 knots for survival and growth, respectively, to determine graphically how well each model captured the mean trend [55]. The probability of reproducing (*p_r_(z)*), and flower production (*f_s_(z)*) had two competing models – the intercept-only, and the log-transformed surface area as fixed effects. The most parsimonious models for each vital rate were selected using AIC. Analysis of the residuals of the growth model showed a gradually declining variance as size increased, and so we fit a model with exponentially decaying variance. As this model had a substantially lower AIC score than any other candidate model, we used it to predict both the mean and variance of the growth distribution. The growth model assumed errors followed a Gaussian distribution. Survival and probability of reproduction models were fit using binomial error distributions. The flower production model was first fit with a Poisson family, but analysis of the deviance showed overdispersion. Thus, we re-fit it with a quasi-Poisson to avoid underestimating standard errors for each coefficient. Results for the model fitting are shown in Table 1 and the model selection process are further described in the Online Supplementary Materials.

**Table 1.**
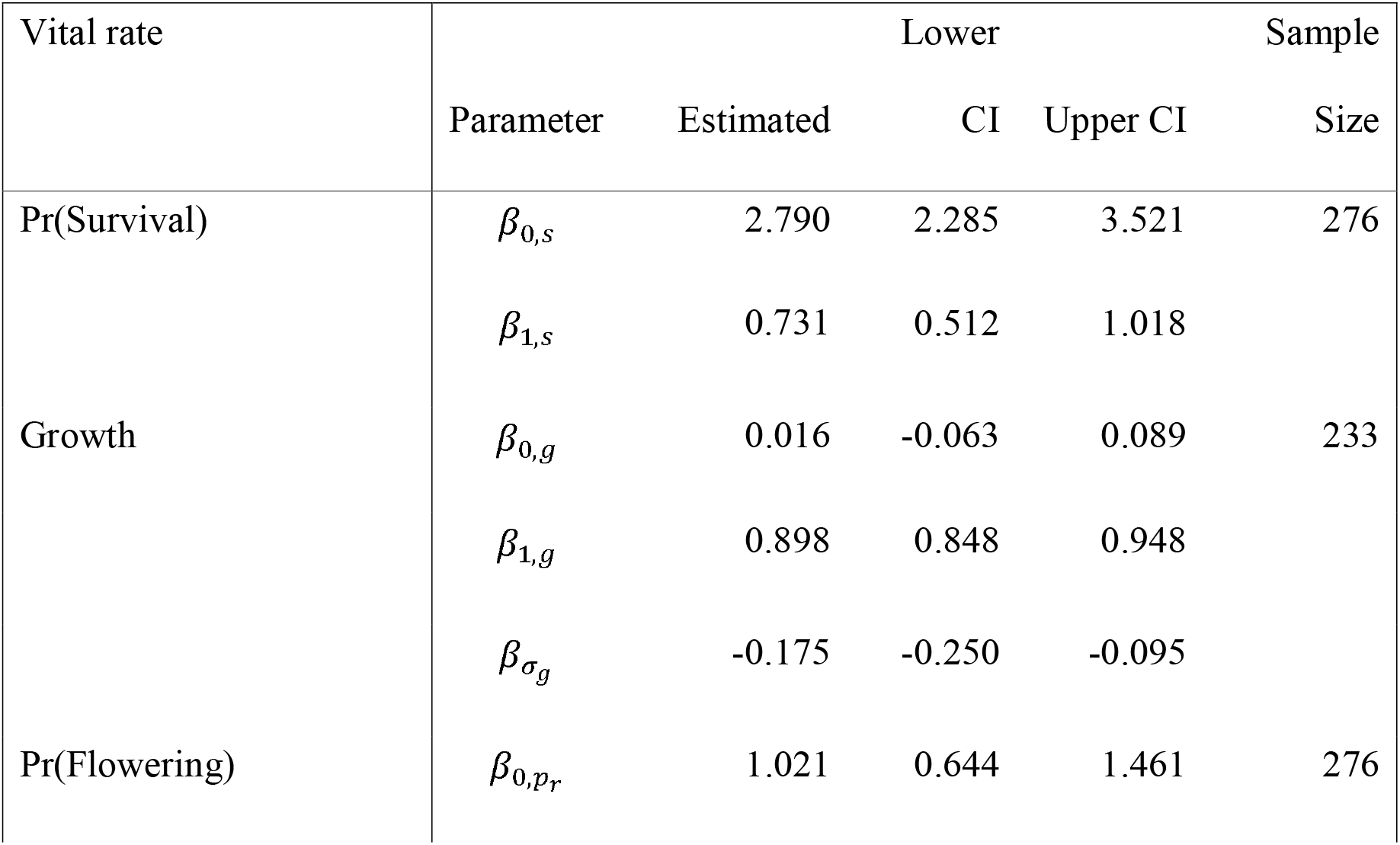
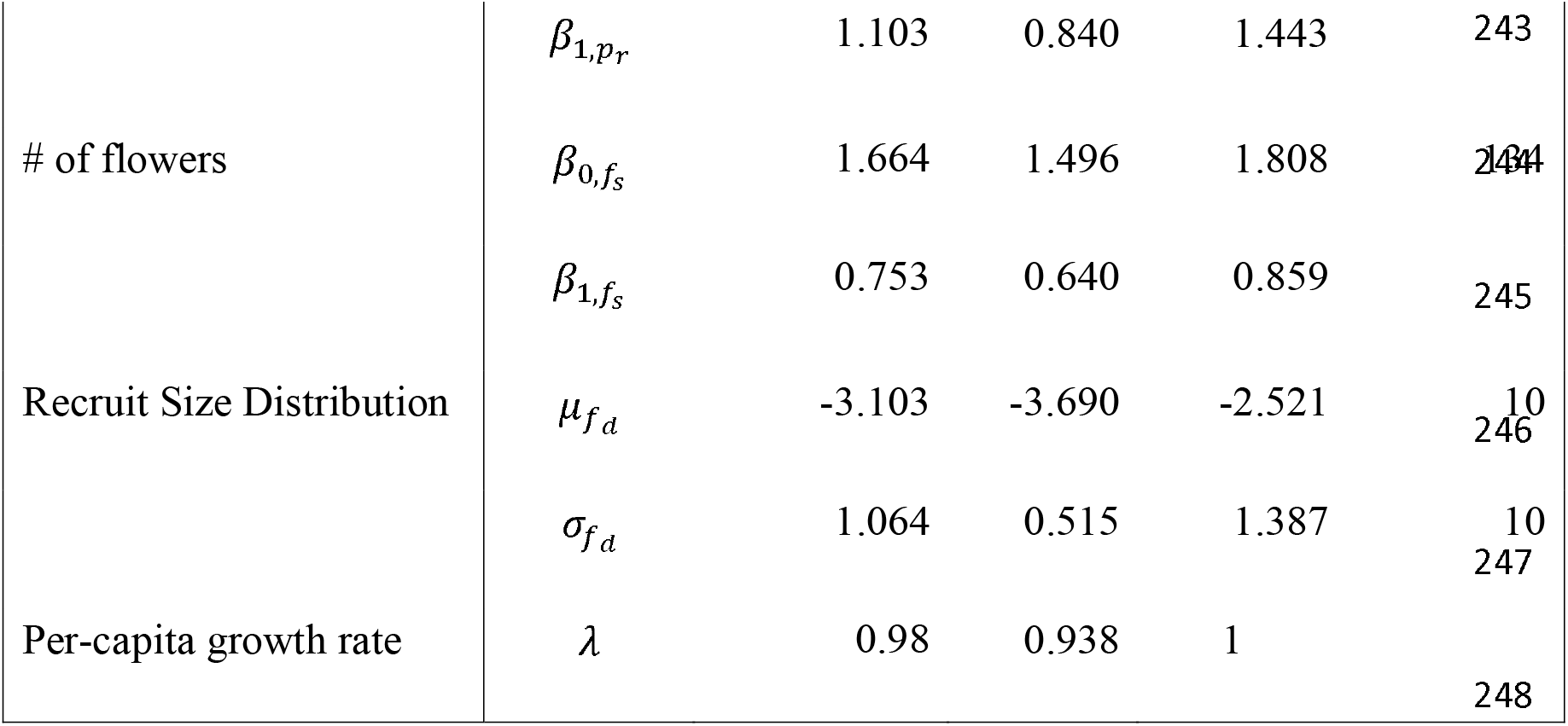
Estimated regression coefficients of vital rates and their 95% confidence intervals. Confidence intervals were obtained by bootstrapping the data set 1000 times and re-fitting the model. Sample size indicates the number of individuals used in parameter estimation.

In addition to the regression models, we used our data to determine the size distribution of new recruits (*f_d_(z’)*) and the number of new recruits in 2019 per flower produced in 2018 (*f_r_*). The recruit size distribution was modelled using a truncated normal distribution, and the rate at which flowers produced new individuals was modeled by dividing the number of new recruits in 2019 by the total number of flowers produced in 2018. *Carpobrotus* flowers take over a year to mature fruits, and thus new recruits found in 2019 are the result of flowers produced prior to 2018. The most parsimonious model for *f_r_* with our data considers flower production to be time invariant (e.g., flowers produced by the population is similar from year to year). We performed a regression parameter-based perturbation analysis to investigate how flower number variation affected our results [25]. We treated each parameter value in the flower number regression model as a Gaussian random variable and drew 1000 estimates from the distributions for each implied by the model estimation procedure. We then used parameter draw values for each parameter to rebuild the model 1000 times and computed the per-capita growth rate again. The per-capita growth rate for the population is not very sensitive to these perturbations (Figure S1.1), and so we feel comfortable that the underlying assumption did not substantially affect our results.

We combined the vital rate functions using an Integral Projection Model (IPM; [20,25]) to estimate the population’s long-term trends. IPMs are similar to matrix projection models (MPMs; [19]) but allow for individuals to be classified by a mixture of discrete and continuous state variables, rather than discrete variables only as in MPMs. These models take the following general form:

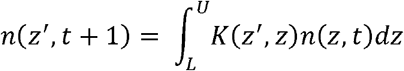

In this equation, *n(z’, t + 1)* is a function describing the size structure of the population at time *t + 1, z’* is the state variable used to describe the population, and the integral of the expression over the domain of *z’* is the total population size. *L* is the lower limit of this domain, and *U* is the upper limit. *K(z’,z)* is a bivariate kernel function that describes transitions to state *z’* given an individual’s initial state, *z*, at time *t*. For this model, log-transformed surface area of individuals is the state variable *z*. *K(z’,z)* is typically comprised of two or three sub-kernels that describe growth of existing individuals conditional on survival (*P*), sexual reproduction (*F*), and asexual reproduction (*C*). These sub-kernels are themselves comprised of functions describing the vital rates that contribute to them, and the functions are based on the regression models described above. Because of *Caprobrotus’s* sprawling growth habit, we were unable to distinguish between growth of surviving individuals, and clone production when clones were contiguously part of the existing ramet (See Discussion). Thus, our model contained only two sub-kernels: *P* and *F.* Substituting these into Eq 1, our model took the form

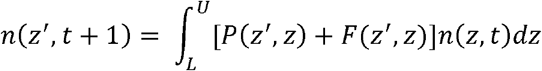

Analytical solutions to these integrals are not available, and so a numerical solution is necessary [25,56]. We solved the equation numerically using the midpoint rule of integration with 100 meshpoints along the domain [*L, U*] to generate a 100 by 100 iteration matrix, *K.* We checked whether using different numbers of meshpoints affected our results by rebuilding the model from 100 to 500 meshpoints at increments of 50 meshpoints each time. The change in percapita growth rate was on the order of 10^-5^, so we retained the model with 100 meshpoints and continued with the analysis. A complete description of the model functions is in Online Supplementary Materials.

Once the IPM was implemented, we then calculated the per-capita growth rate as the dominant eigenvalue (*λ*) of the iteration matrix, K [25]. We also computed the sensitivity and elasticity functions of *λ* at the kernel and sub-kernel level to understand the relative contributions of survival/growth transitions and reproductive transitions to the population growth rate [25]. To quantify uncertainty in our data, we resampled our data with replacement 1,000 times and refit all vital rates using the same functional forms. We constructed IPMs for each iteration and computed per-capita growth rates, sensitivities, and elasticities. Thus, all estimates reported here include our point estimate and the 95% confidence intervals from the bootstrapping procedure.

All the code for the statistical analysis is available at the following link https://github.com/levisc8/carpobrotus_IPMs/tree/master/Ana_Israel_IPM.

## Results

Vital rate regression using log-transformed surface area of individuals of *C. edulis* as a fixed effect yielded good fits for all of our vital rates and generally lower AIC scores than intercept-only, quadratic fits, or GAMS (Figure 2, Table 1, Online Supplementary Materials). GAMs and polynomial fits did not substantially improve the overall model fit or substantially lower the AIC score for each vital rate, even with the increased complexity of the model itself. Thus, our IPM only includes models with a single fixed effect –log-transformed surface area– as explanatory variables (Online Supplementary Materials).

**Figure 2:**
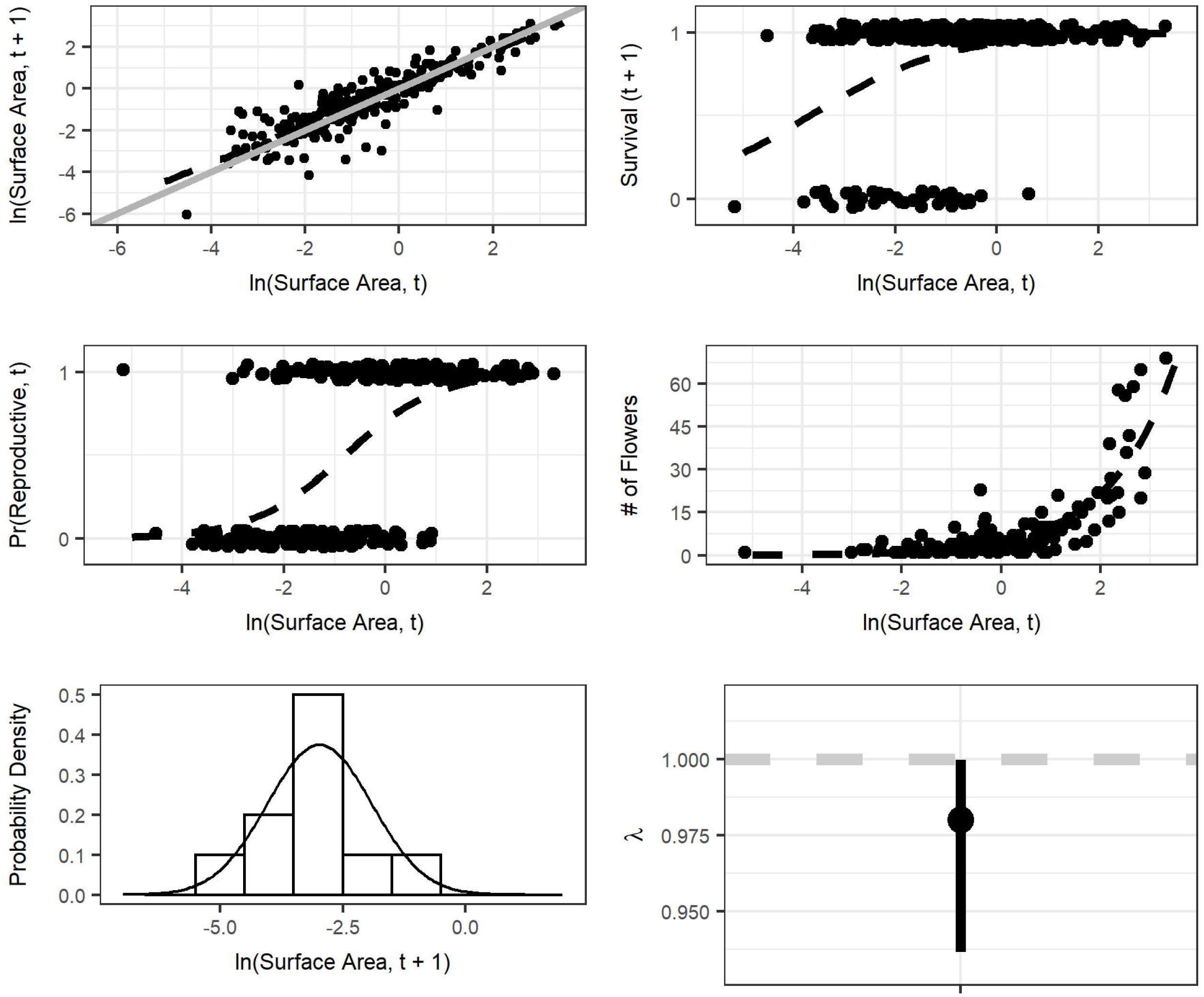
Population vital rate results for growth (a), survival (b), probability of flowering (c), flower production (d), recruit size distribution (e), and per-capita growth rates (f). The dotted black lines in a-c, e and the solid black line in d are fitted relationships from the models described in Methods. In (a), the solid grey lines show a 1:1 relationship (e.g. individual level stasis), and the dotted grey line in f denotes a per-capita growth rate of 1 (e.g. population level stasis). The range line around the point estimate of the per-capita growth rate shows the 95% confidence interval derived from the boot strapping procedure.

The emerging estimate of population growth rate was *λ* = 0.98 (Lower 95% CI = 0.938, Upper 95% CI = 1). These estimates indicate that the population would not significantly depart from demographic stability on the long-term under conditions of environmental constancy and density independence (Figure 2). The IPM’s sensitivity surface indicates population growth rates are most sensitive to small individuals growing large rapidly, and the elasticity analysis indicates that survival/growth transitions are far more important to *λ* than reproduction (Figure 3).

**Figure 3:**
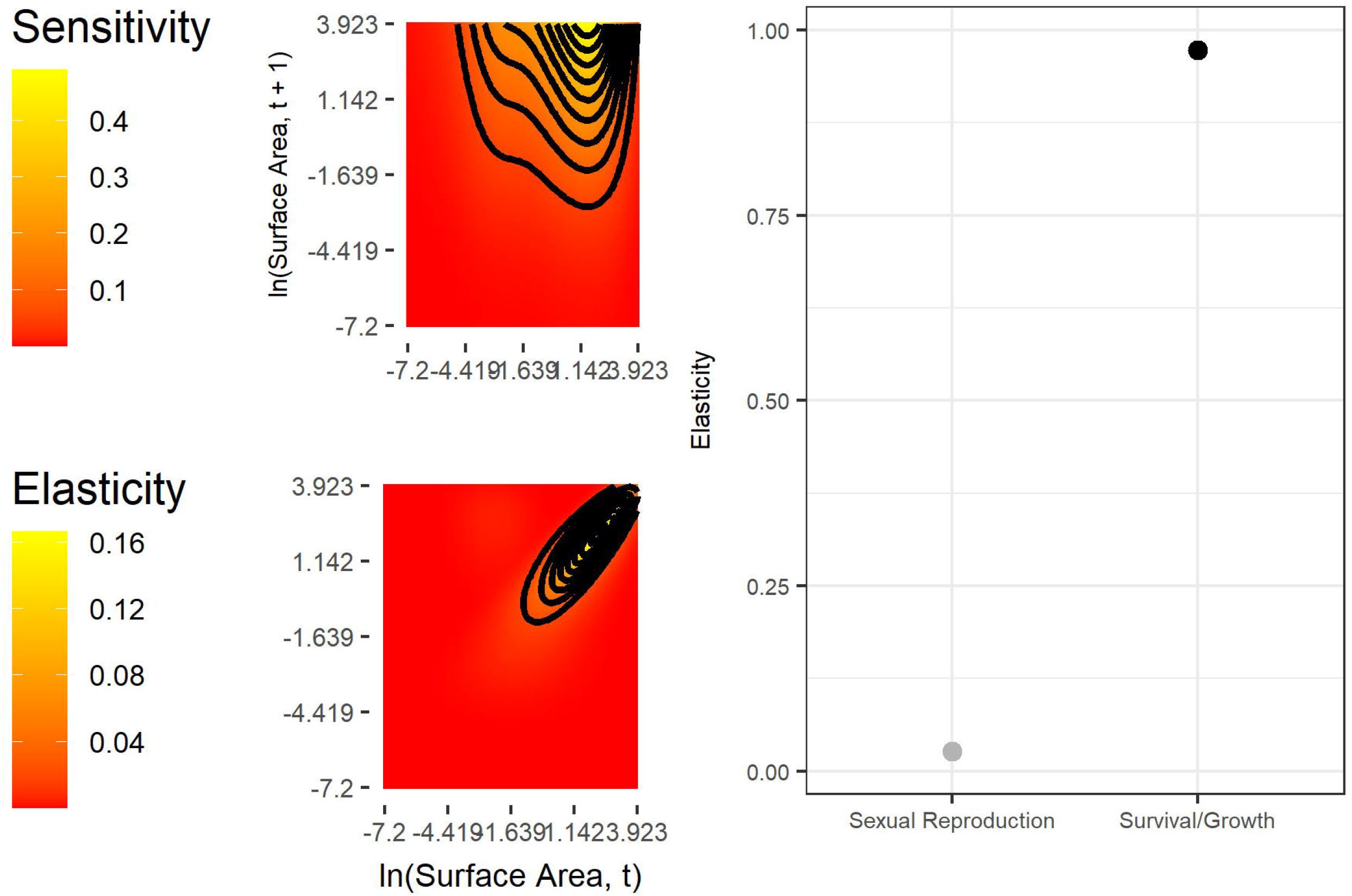
Left: Sensitivity and elasticity kernels for the complete model. Right: Elasticities for each sub-kernel, indicating relative contributions of each to the per-capita growth rate. The range line each point estimate of the sub-kernel level elasticity shows the 95% confidence interval derived from the boot strapping procedure

## Discussion

Demographic models are ideal to understand population processes and stages in the life cycle of species that require urgent management measures, such as the harmful invaders of the *Carpobrotus* genus. These are species of conservation and economic concern that thus far remained unexplored demographically, these results providing the first stage-based population projection model in the entire Aizoaceae family. Such structured population models of invasive plant species provide a tool for generating comprehensive fitness estimates and quantifying relative importance of vital rates (e.g. survival, growth). Our Integral Projection Model results show a population with a growth rate overlapping with the demographic stability boundary of 1, indicating that this population is neither declining nor expanding if the environmental conditions under which it was examined do not change. There are a few reasons that might explain why *Carpobrotus,* which is known to be highly invasive in other places [6,26–28], might have stable growth rate at our site in Israel. First, our Havatselet population might be an old population that has reached an ‘equilibrium’ and is now limited by space [24,57,58]. The population grows on a cliff side and may be difficult for it to spread to unoccupied spots. Individual plants may produce fruits and seeds that fall directly in the Mediterranean Sea and their roots may have difficulty finding support on the bare vertical cliff. Second, the Israelian climate might not be ideal for *Carpobrotus.* The Cape Region, where these plants are native, has a cold semi-arid climate [59], with average monthly temperatures reaching 21.60□ [6]. Our Havatselet site has a warm Mediterranean climate [59] with higher temperatures, reaching a mean of 27.0□. Summers in Israel are much drier, often with no precipitation, compared to the minimum of 30 mm of rain the summer months of Cape Region receives [60]. Third, it is possible that stable population growth rates are typical for *Carpobrotus* and that its invasive character is rather due to its ability to spread regionally, either naturally or through human-mediated introduction. *Carpobrotus* was introduced multiple times all over the world for ornamental and soil stabilization purposes and continues to be planted by people [61].

We find that the population growth rate of our *Carpobrotus* population is highly sensitive to changes in survival and growth of plants and less sensitive to reproduction. The sensitivity analysis indicates that rapid growth of plants that are already at a moderate size would have dramatic effects on *»* We did not observe any individuals in our study having such dramatic changes in size and it might not be biologically possible for *Carpobrotus.* Our result that elasticity of *λ* is high for survival and low for sexual reproduction is expected. Across the plant and animal kingdoms, elasticity of *λ* to survival is known to increase with life span [24,62–65], suggesting that persistence of healthy adults is critical to maintaining populations of long-lived perennial plants.

While the population growth rate *λ* we obtained with our model suggests that the population is long-term demographically viable, it is important to note that we have only analyzed one annual transition. It is possible that rare and favorable environmental conditions in other years could allow this population to undergo accelerated population growth. However, even if the *Carpobrotus* population remains stable, it still greatly hinders any attempt of native species reintroduction or management, due to its direct and indirect effects on the environment as an invasive transformer (e.g., allelochemical activity that disrupt seed germination and root growth of native plants; [15,16]). For these reasons, the *Carpobrotus* population reduction is advisable. Further monitoring is required to answer the question of how many individuals would need to be removed, but other studies have shown that plant populations can respond in a non-linear manner to large perturbations (e.g., [66]).

Our elasticity analysis indicates that large adults should be targeted for removal. Unfortunately, most of these large plants are on the face of the cliff, and their removal would be challenging, time-consuming, and expensive. Mechanical control is not possible, and the chemical methods are not ideal. Even if plants could be reached and herbicide could be applied, these habitats are windy, and the herbicides could get in the ocean or affect native flora [6]. Biological control agents that reduce survivorship are under consideration. Two accidentally introduced soft-scale insects, *Pulvinariella mesembryanthemi* and *Pulvinaria delottoi,* caused severe damage to *C. edulis* plantings in California in the 1970s. At the time, *C. edulis* was still regarded as a desirable plant so the California highway Department introduced natural enemies to control the insects. More recently, scale insects have been observed to cause mortality of *Carpobrotus* in California, suggesting these might have the requisite efficacy [6,67]. Since 2015, the control efficiency of the generalist pathogenic fungus *Sclerotinia sclerotiorum* and the soft scale insect *Pulvinariella mesembryanthemi* are being assessed, but definitive results are yet to be seen [6]. Finally, resident generalized grazers, such as deer and rabbits can reduce the establishment and growth of *Carpobrotus,* but also might disperse seeds and create disturbances and allow for seedling establishment [6,67].

We suggest, as other researchers have, that prevention is critical to curb the regional spread of *Carpobrotus.* To date, *C. edulis* and *C. acinaciformis* are still available to the public and are allowed to be planted in urban gardens that are not adjacent to natural areas and that are outside of the coastal plains [48]. We recommend containment of the species in the regions it has already invaded and prevention of its human-mediated spread. Currently, there is not adequate legislation in most countries for such prevention. Some European countries have included *C. edulis* in invasive species lists and thus forbid its release in the environment. To our knowledge, Israeli legislation does not include rules for the introduction and control of invasive species [6,68].

We were successful in using the drone for creating the maps of the *Carpobrotus* population and later use those maps for evaluating vital rates and parameterizing an IPM. This is another potential ecologically use case for a rapidly developing technology used in both animal [45,69,70] and plant population studies [42,71,72]. While in our case, the drone successfully generated maps and subsequent estimates of plant size, we acknowledge that they have their limitations. Georeferencing can prove challenging in environments without many permanent features to mark across multiple samples. If maps are not well aligned, re-identifying individuals from the previous year can be quite challenging. Advances in RTK and PPK GPS technology have greatly reduced this challenge, though these can still be quite pricey relative to typical conservation budgets. Furthermore, size estimates can vary greatly when the absolute distance between the ground and the camera varies across the orthomosaic scene, necessitating the placement of ground truth targets in areas that may be difficult to physically reach. Additionally, dense vegetation mats can cause confound the orthomosaicing procedure. Even when it is successful, it may be difficult to identify individual plants. Therefore, we recommend that vegetation density and height be considered when choosing the mapping method.

In conclusion, we confirm that the size of different *Carpobrotus* individuals is a reliable predictor for vital rates, and that the demography of *Carpobrotus* is typical of a long-lived polycarpic plant. We demonstrated that drones can successfully allow for demographic data collection for plant species that were previously inaccessible to researchers. Thus, using this technology in future demography research shows great promise for filling global data gaps. Our results also have consequences for possible *Carpobrotus* control actions, as we emphasize the need to target survival and growth of large plants.

