## Appendix S1 for "Demographic analysis of Israeli *Carpobrotus* populations: management strategies and future directions"

### IPM Functions

1. $n(z', t+1) = \int_L^U K(z', z)n(z, t)dz$
2. $K(z', z) = P(z', z) + F(z', z)$
3. $P(z', z) = s(z) * g(z', z)$
4. $F(z', z) = p_r(z) * f_s(z) * f_d(z') * f_r$

### P(z',z)

5. $Logit^{-1}(s(z)) = \beta_{0,s} + \beta_{1,s} * z$
6. $g(z', z) \sim Norm(\mu_g, \sigma_g)$
7. $\mu_g = \beta_{0,g} + \beta_{1,g} * z$
8. $\sigma_g = \sqrt{e^{2*\beta_{\sigma_g}*z}}$

### F(z', z)

9. $Logit^{-1}(p_r(z)) = \beta_{0,p_r} + \beta_{1,p_r} * z$
10. $ln(f_s(z)) = \beta_{0,f_s} + \beta_{1,f_s} * z$
11. $f_d(z') \sim Norm(\mu_{f_d}, \sigma_{f_d})$

##### Implementation rules

Integration rule: midpoint

L = -7.256

U = 3.979

### meshpoints = 100

### Drone results

The drone surveys from 2018 required 6 batteries over the course of 2 days due to a limited battery supply. Ambient temperature was approximately 24 degrees and there were no clouds present during either sampling period. In total, 1720 images were taken of the population from a height of ~9m above ground level. Due to difficulties in following the terrain perfectly, pixel sizes varied, though were on average 0.26cm/pixel. Pix4DMapper successfully geolocated and matched 1695/1720 images across 6 blocks, with a median of 10005.9 matches/image and a 1.83% difference in initial and optimized camera parameters.

The 2019 survey was conducted on a single day under similar conditions to 2018. 1024 images were collected 10-15m above ground level to increase the variability within each image and prevent Pix4DMapper from generating blocks during orthomosaic generation. 1021/1024 images were successfully calibrated in a single block with a mean resolution of 0.45cm/pixel. Despite the drop in resolution, flowers and plants were still quite easy to identify in the resulting orthomosaic. Images had a median of 10219.3 matches, and there was a 0.35% difference between initial and optimized camera parameters.

At each sampling, squares of known size were also recored in each image and had polygons drawn around them (each target measured 0.375m X 0.315m). Thus, we were able to compute the ratio of observed surface areas to true surface areas for the orthomosaic. Observed plant surface areas were standardized using the ratio of the true size to the observed sizes.

### Model fitting results

Table S1.1: Coefficients, standard errors, test statistics, and p-values for each of the vital rate regression used in the IPM to generate point estimates of lambda, sensitivity, and elasticity functions.

| Vital Rate | Parameter | Estimate | Std. Error | Test Statistic | P-value |
|---|---|---|---|---|---|
| Growth | Intercept | 0.02862 | 0.0397 | 0.721 | 4.72e-01 |
| | Slope | 0.89849 | 0.0252 | 35.595 | 9.59e-96 |
| | Sigma Exponent | -0.1754 | | | |
| Survival | Intercept | 2.6963 | 0.313 | 8.60 | 7.84e-18 |
| | Slope | 0.7312 | 0.144 | 5.09 | 3.51e-07 |
| Pr(Flowering) | Intercept | 0.8801 | 0.192 | 4.59 | 4.33e-06 |
| | Slope | 1.1028 | 0.136 | 8.09 | 6.20e-16 |
| Flower Production | Intercept | 1.568 | 0.0453 | 34.7 | 3.47e-263 |
| | Slope | 0.753 | 0.0238 | 31.6 | 1.07e-219 |
| Recruit Size | Mu | -2.975 | | | |
| | Sigma | 1.064 | | | |

Table S1.2: AIC model selection tables for candidate growth models.

| Model | Type | df | AIC |
|---|---|---|---|
| Growth | Intercept Only | 2 | 863.1472 |
| Growth | Linear Term + Intercept | 3 | 483.1242 |
| Growth | Quadratic Term + Intercept | 4 | 484.9146 |
| Growth | GAM | 3 | 483.1242 |
| Growth | Exponential Variance | 4 | 459.3743 |

Table S1.3: AIC model selection tables for candidate survival models.

| Model | Type | df | AIC |
|---|---|---|---|
| Survival | Intercept Only | 1.000 | 240.8141 |
| Survival | Linear Term + Intercept | 2.000 | 208.5736 |
| Survival | Quadratic Term + Intercept | 3.000 | 209.2357 |
| Survival | GAM | 2.282 | 208.5197 |

Table S1.4: AIC model selection tables for candidate probability of flowering models.

| Model | Type | df | AIC |
|---|---|---|---|
| Probability of Reproduction | Intercept Only | 1 | 384.3853 |
| Probability of Reproduction | Linear Term + Intercept | 2 | 273.8571 |

Table S1.5: AIC model selection tables for candidate flower production models.

| Model | Type | df | AIC |
|---|---|---|---|
| Flower Production | Intercept Only | 1 | 2039.4341 |
| Flower Production | Linear Term + Intercept | 2 | 890.0934 |

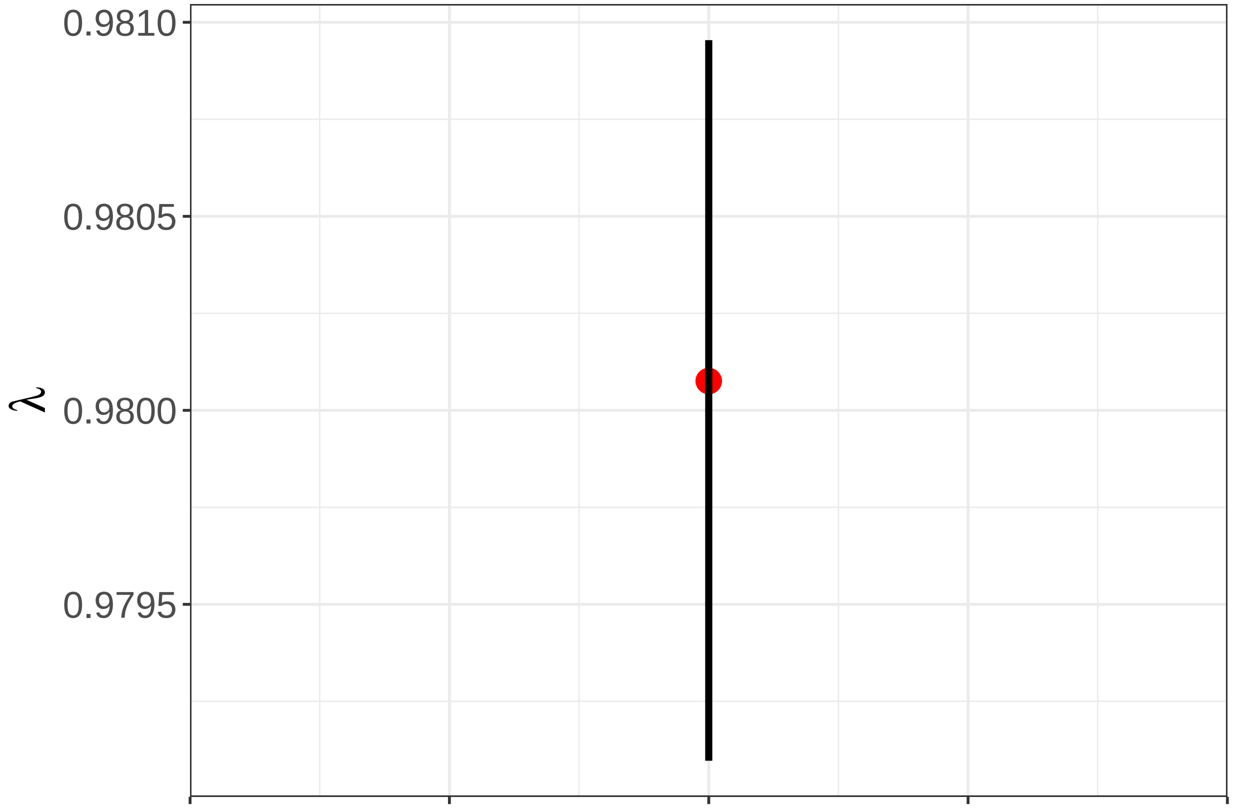


Figure 1: Results for the flower number regression parameter perturbation analysis. The red point is the per-capita growth rate estimated from the regression parameters. The black line shows the upper and lower 95% confidence intervals derived from the draws that perturbed each flower number regression parameter. The range of values is quite small, and our qualitative results are not affected by the assumption of little temporal variation in flower production.
